# Comparative Hox genes expression within the dimorphic annelid *Streblospio benedicti* reveals patterning variation during development

**DOI:** 10.1101/2023.12.20.572624

**Authors:** Jose Maria Aguilar-Camacho, Nathan D. Harry, Christina Zakas

## Abstract

Hox genes are transcriptional regulators that elicit cell positional identity along the anterior-posterior region of the body plan across different lineages of Metazoan. Comparison of Hox gene expression across distinct species reveals their evolutionary conservation, however their gains and losses in different lineages can correlate with body plan modifications and morphological novelty. We compare the expression of eleven Hox genes found within *Streblospio benedicti,* a marine annelid that produces two types of offspring with distinct developmental and morphological features. For these two distinct larval types, we compare Hox gene expression through ontogeny using HCR (hybridization chain reaction) probes for *in-situ* hybridization and RNA-seq data. We find that Hox gene expression patterning for both types is typically similar at equivalent developmental stages. However, some Hox genes have spatial or temporal differences between the larval types that are associated with morphological and life-history differences. This is the first comparison of developmental divergence in Hox genes expression within a single species and these changes reveal how body plan differences may arise in larval evolution.

## INTRODUCTION

Hox genes are a classic example of conserved transcriptional regulators that assign segment identity along the anterior-posterior axis in all Metazoans (Mallo and Alonso, 2013; Hombria et al., 2021), even cell positional identity (Yu et al., 1995; Afzal and Krumlauf, 2022). Therefore, they have received much attention when it comes to body plan evolution (Mallo and Alonso, 2013; Afzal and Krumlauf, 2022). But the extent that differences in Hox gene expression (in both space and time) contribute to developmental and morphological differences depends on many phylogenetic and developmental factors (Fröbius and Funch, 2017; Martin-Zamora et al., 2023). Here we use an annelid with two distinct developmental pathways to compare how Hox gene expression shapes developmental and morphological differences.

While the conserved Hox gene expression pattern and synteny across distant taxa are renowned, there are many examples where changes in Hox gene expression have led to evolutionary diversification (Lee et al., 2003; Monteiro and Ferrier, 2006). In selected model organisms where Hox gene expression has been investigated, the genes appear in ‘clusters’ and are expressed in domains along the anterior-posterior axis at different developmental stages (Shankland and Seaver, 2000; Fröbius and Funch, 2017; Krumlauf, 2018). The expression patterns follow the same sequential order as their genome location, displaying spatial and temporal colinearity (Denans et al., 2015; Mallo, 2022).

There are, of course, exceptions to these rules. Hox genes are not always arranged in clusters; they may be on a single chromosome but spatially distinct (e.g., the mollusk *Crassostrea gigas* and the marine annelid worm *Capitella teleta* have a distantly located anterior Hox cluster; Fröbius et al., 2008; Li et al., 2020). They could be in different chromosomes (e.g., in the spot octopus *Octopus binocularis* and the flatworm *Schmidtea mediterranea*; Albertin et al., 2015; Currie et al., 2016). Also, Hox gene expression may deviate from spatial and temporal collinearity. For example, most of the Hox genes from the tunicate *Oikopleura dioica* are expressed in the tail and not in the anterior region at the larval stage; and in the brachiopod *Terebratalia transversa* Hox genes are not expressed collinearly at distinct developmental stages (Seo et al., 2004; Monteiro and Ferrier, 2006; Schiemann et al., 2017). Hox genes may also take on different functions from the canonical expectation of specifying anterior-posterior segment identity. Hox genes can be co-opted or recruited to other functions that are lineage specific (For Sprialian examples, *Post1* is expressed in the developing buccal lappets and in the developing light organ in the larva of the squid *Euprymma scolopes;* and this same orthologue is expressed in the swimming chaetal sacs in the notochaete larva of the marine worm *Nereis virens* (Lee et al., 2003; Kulakova et al., 2007).

To what extent do differences in Hox gene expression— in either time or space— lead to life-history and developmental changes? Studies of Hox gene expression patterns in annelids have shown a conserved expression pattern across related species, despite distinct larval life-histories (Kulakova et al., 2007; Fröbius et al., 2008; Bakalenko et al., 2013). However, Hox gene co-option is related to morphological novelties that are lineage-specific (Kulakova et al., 2007; Fröbius et al., 2008; Bakalenko et al., 2013).

While many comparisons have been made across species, we take advantage of a model with an intraspecific developmental dimorphism to determine whether Hox gene expression differs based on development mode. This is essentially the opposite approach of comparing across distant lineages, instead looking at the closest possible evolutionary distance to identify the effects of subtle changes. Here we ask if Hox genes drive developmental divergence within a species. We use the estuarine annelid *Streblospio benedicti* that produces two types of offspring with distinct embryological and morphological features (reviewed in Zakas, 2022). In this species, females brood embryos in a dorsal brood pouch before releasing them as swimming larvae. There are two types of females: They either produce many small eggs that develop as obligately-feeding planktotrophic larvae, or a few large eggs that develop as non-feeding lecithotrophic larvae. Planktotrophic larvae spend 2-3 weeks feeding in the plankton before becoming competent to metamorphosis, while lecithotrophic larvae can settle to the benthos within a day of release. In addition to egg and larval size, there are distinct morphological differences between the swimming larval types. For example, planktotrophic larvae have swimming chaetae which are absent in the equivalent swimming lecithotrophic larvae. (Fig. 1). By using this model, we compare Hox gene expression differences across divergent life-history modes while controlling for interspecific evolutionary differences. By determining Hox gene expression patterns at distinct developmental stages in the two developmental modes we add to the growing understanding of annelid Hox gene evolution.

**Fig. 1.**
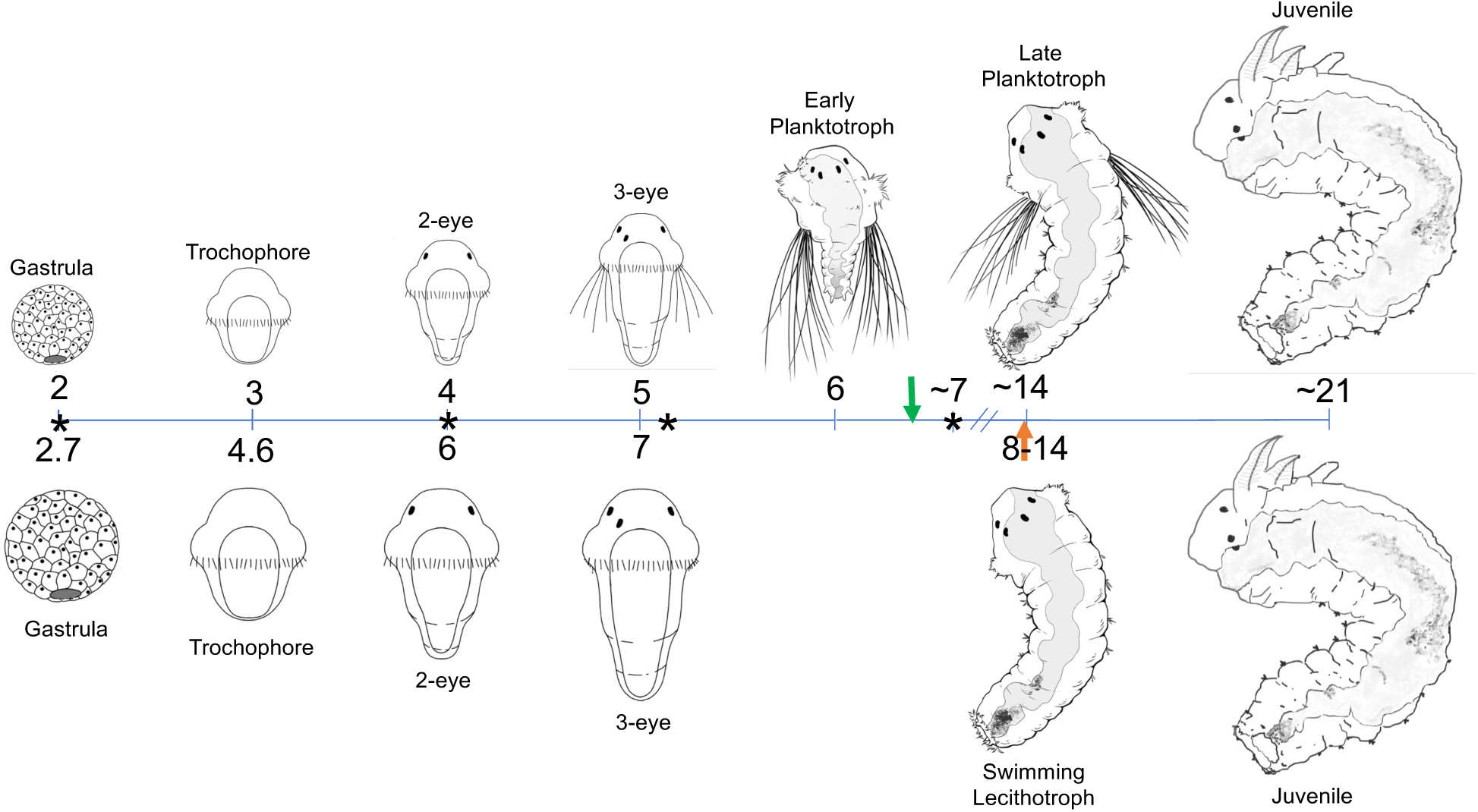
Timeline of larval development in days post fertilization. Both developmental types reach the same embryonic stages. Time is in days post fertilization (dpf). Lecithotrophic embryos are delayed in developmental timing compared to planktotrophic larvae at the equivalent stage. Planktotrophic larvae have an extra swimming stage compared to the lecithotrophic larvae. Both types are morphologically indistinct by the early juvenile stage. Green and orange arrows indicate the approximate time when larvae are released from their mother. Asterisks indicate the RNAseq stages: gastrula, 2-eye, swimming larvae, and 1-week larvae.

## RESULTS

### Larval staging and developmental differences

The distinct larval and life-history dimorphism in *S. benedicti* is well characterized (reviewed in Zakas, 2022). While both larval morphs reach equivalent stages through unequal spiral cleavage, they differ in size and the absolute development time to each stage (Fig. 1). Despite this difference, the blastula, gastrula, and trochophore stages (where there is a distinct prototroch ciliary band and pear-shaped larva) are remarkably similar. Small planktotrophic larvae develop faster than their larger, yolky lecithotrophic counterparts (∼ 8x difference by volume), which is expected as the larger cells take longer to divide (McCain, 2008).

We identified equivalent embryological and larval stages based on their conserved developmental patterns (Fig. 1). For this study we investigated gastrula and trochophore stages for early gene expression. We termed the next stage as the ‘**2-eye’** stage. At this point the larvae develop two red eyes simultaneously as the first anterior body segment of the trunk appears. Aside from size and yolk content, planktotrophic and lecithotrophic larvae appear similar at this stage.

Notable morphological differences arise at the ‘**3-eye’** stage. Larvae add an additional anterior-lateral eye on one side (usually the left) a day later. (While eye number is usually obvious, we also control our staging by development time, using larvae 24 hours post 2-eye stage). The posterior region consists of a growth zone with two or four segments below the head. One major difference is that only the planktotrophic larvae grow ‘swimming chaetae’ which originate from paired, lateral chaetal sacs on the first body segments. The lecithotrophic offspring never produce these swimming chaetae, although they do have equivalent first chaetal sacs. Swimming chaetae, or “provisional chaetae” are morphologically distinct from the smaller body chaetae found in the lateral, paired, parapodia of each segment.

The next developmental stage is when the larvae are typically released from the mother’s brood pouch and most of the characteristic life-history differences occur. Larvae have four red eyes (two pairs) at this stage. These **‘early-planktotrophic**’ larvae can swim and ingest food as they have a functional mouth; a through-gut, a short trunk, and pronounced anal cirri on the tail. The swimming chaetae extend the full body length (Pernet and McArthur, 2006). The ‘**late-planktotrophic**’ larval stage is approximately 10 days later, when the larvae have added additional posterior segments and feed, but they have not shed their swimming chaetae. We selected larvae that have ∼12-13 segments for analysis, but planktotrophic larvae can take 2-4 weeks in the plankton at this stage before metamorphosing into juveniles. This ‘late-planktotrophic’ stage is morphologically like the ‘**swimming lecithotrophic’** larval stage, with the notable exception of lecithotrophic larvae lacking swimming chaetae or pronounced anal cirri. The ‘swimming lecithotrophic’ larvae also have ∼12-13 segments. Notably, the lecithotrophic larvae are released from their mother at this stage and only take ∼1 day before metamorphosing. Therefore, females of each type brood their larvae for different periods of time: planktotrophic larvae are released at the early-planktotrophic stage around 7 days post fertilization (dpf), while lecithotrophic larvae are released when they are competent to metamorphose, closer to 12-14 dpf. The time they spend as pelagic larvae is drastically different. However, at this late larval stage— just prior to metamorphosis— the two larval types are comparable in body plan and size. Both developmental types are sequentially adding segments to the posterior growth zone, and both appear competent to metamorphose.

At the ‘**juvenile stage’** the two developmental modes converge in body plan and are morphologically indistinct. The juvenile has a peristomium and prostomium and a segmented body with ∼13 segments (below the peristomium). Two palps and two branchiae arise from the peristomium. In each segment, the notochaetae and neurochaetae are present. The late-planktotrophic larvae shed their swimming chaeta when undergoing metamorphosis to the early juvenile stage. It takes approximately four weeks for the early juvenile to become a reproductive adult (Zakas, 2022).

### Identification of the Hox genes

We identified eleven Hox genes in *S. benedicti* using a homologous gene approach: We assembled a transcriptome from RNAseq data of each developmental stage in the two developmental modes. We blasted known Hox genes from other spiralian species against our transcriptome to identify homologs in *S. benedicti*. We verified the best hits were genic regions with open reading frames.

The 11 Hox genes are located on chromosome 7 of the *S. benedicti* genome, with an anterior cluster (*Lab, Pb, Hox3, Dfd, Scr, Lox5*, *Ant* and *Lox4*) that spans ∼463 000 kbp and a separate posterior cluster (*Lox2, Post2* and *Post1*) further away on chromosome 7 (Zakas et al., 2022; Figs. 2, Table S3). We constructed the molecular phylogenetic tree of the Hox genes from aligned amino acid sequences across spiralian species. We find *S. benedicti* Hox genes cluster within clades of ortholog spiralian Hox genes (Fig. 3, S1). This indicates that there are no duplication events for the Hox genes in the genome and gives high confidence that the Hox gene assignments are correct.

**Fig. 2.**
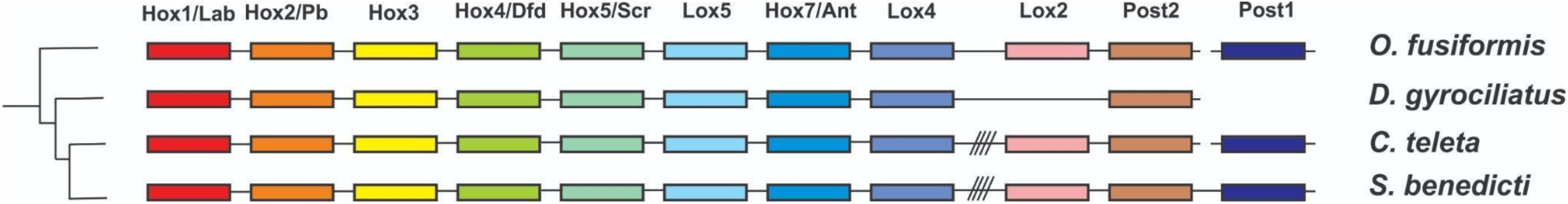
Schematic representation of the Hox genes in *S. benedicti* and other polychaete species. Comparisons with *O. fusiformis* (Martin-Zamora et al., 2023)*, D. gyrociliatus* (Martin-Duran et al., 2021) and *C. teleta* (Fröbius et al., 2008).

**Fig. 3.**
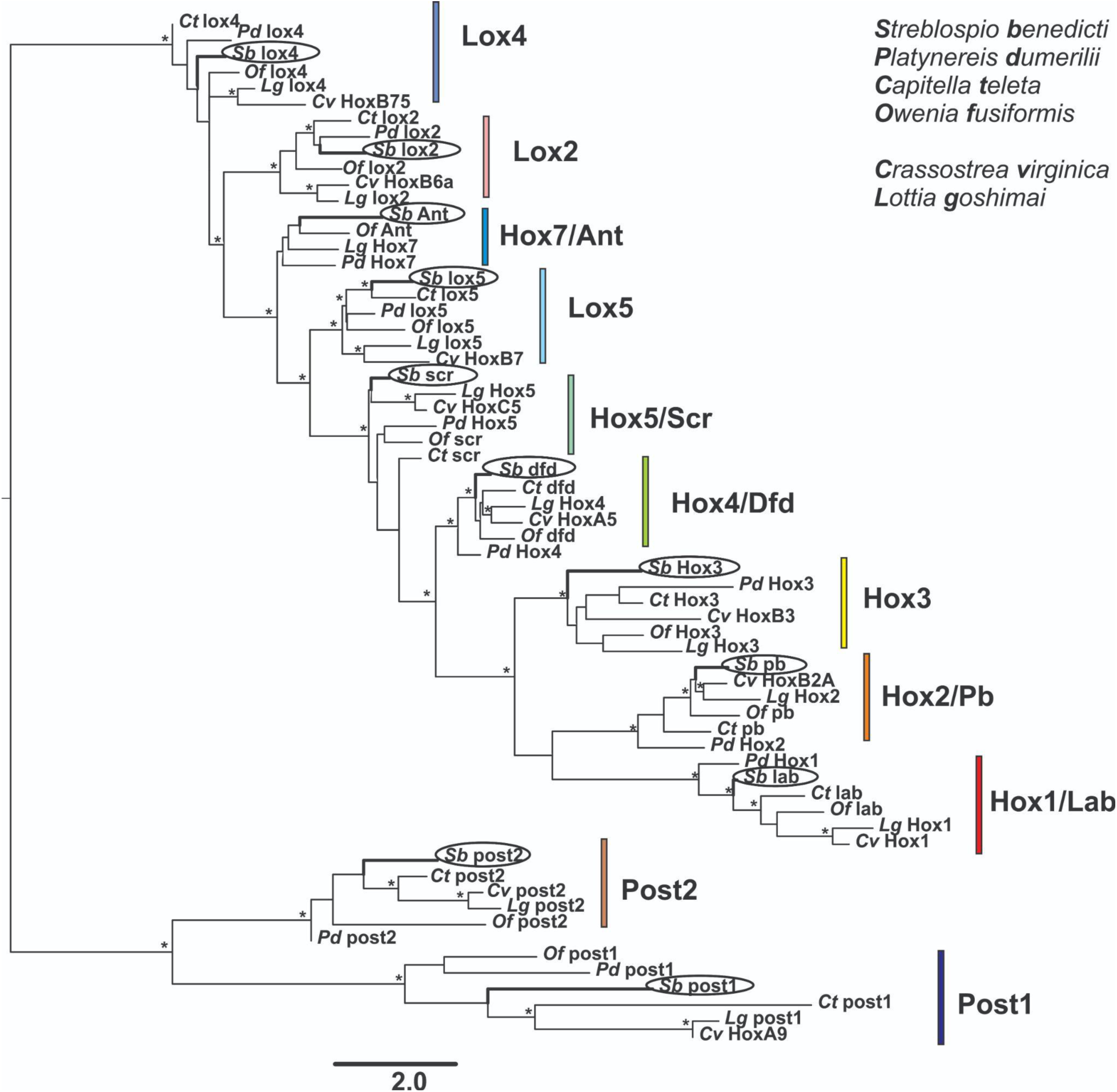
Molecular phylogenetic tree of spiralian Hox genes and orthology assignment for *S. benedicti* Hox genes. The tree was constructed with the IQ-Tree software. Asterisks above branches denote a support value >70% for the three different methods calculated in IQTREE software.

### Hox gene expression patterns during development

To spatially and temporally visualize Hox gene expression, we developed probes for each of the eleven Hox genes using *in situ* hybridization chain-reaction (HCR; Kuehn et al., 2022). To investigate body plan and developmental differences, we first identify the onset of Hox gene expression, as it could be different between the two developmental modes. At the gastrula and trochophore stage, no expression was detected for any of the eleven Hox genes in either developmental mode. The 2-eye stage is the earliest stage where we could visualize expression of any Hox gene (Figs. 4A, 4B,5,6). ***Lab*** is expressed in ventral endodermal cells in segments 2 and 3 in the late-planktotrophic larvae, swimming lecithotrophic larvae, planktotrophic and lecithotrophic juvenile. *Lab* is expressed in the late-planktotrophic larvae and not in the early-planktotrophic larvae. ***Pb*** is expressed in endodermal cells of the swimming chaetal sacs in the early-planktotrophic larvae and cells forming irregular circular spots in the latero/ventral part (segments 3-7) in the swimming lecithotrophic larvae (Figs. 4B, S2). *Pb* has different expression patterns at the swimming larval stage (early-planktotrophic larvae and swimming lecithotrophic larvae) in the two developmental modes. *Pb* is expressed in the early-planktotrophic larvae and not in the late-planktotrophic larvae. ***Hox3*** is expressed in endodermal cells at the posterior end in the 2-eye and 3-eye stages in the two developmental modes, at the mid and lower trunk in the early-planktotrophic larvae, and in segments 8-13 in the swimming lecithotrophic larvae, late-planktotrophic larvae, planktotrophic juvenile, and lecithotrophic juvenile. ***Dfd*** is expressed in cells of the mid-trunk in the early-planktotrophic larvae and in segments 4-8 in the swimming lecithotrophic larvae. *Dfd* is expressed in the early-planktotrophic larvae and not in the late-planktotrophic larvae. ***Scr*** is expressed in endodermal cells in some segments of the mid-trunk in the early-planktotrophic larvae, in segments 4-8 in the swimming lecithotrophic larvae; in segments 4-9 in the late-planktotrophic, planktotrophic juvenile and lecithotrophic juvenile. ***Lox5*** is expressed in some cells of the segments of the lower trunk in the 3-eye stage in the two developmental modes and in the early-planktotrophic larvae; in segments 8-12 in the late-planktotrophic larvae and in segments 5-11 in the swimming lecithotrophic larvae. ***Ant*** is expressed in endodermal cells in one or two segments at the lower trunk in the early-planktotrophic larvae; in segments 6-11 in the swimming lecithotrophic larvae; in segments 7-10 in the late-planktotrophic larvae, planktotrophic juvenile, and lecithotrophic juvenile. ***Lox4*** is expressed in cells in segments 6-12 in the swimming lecithotrophic larvae, planktotrophic juvenile, and lecithotrophic juvenile. *Lox4* is expressed earlier in the lecithotrophic larvae than in the planktotrophic larvae. ***Post2*** and ***Lox2*** No expression was detected for any stages in the two developmental modes (Figs 4A, 4B). ***Post1*** is expressed in endodermal cells at the lateral sides of the mid part in the 2-eye stage of the planktotrophic larvae; endodermal cells forming circular spots at the lateral sides of the mid and lower trunk in the 3-eye stage of the lecithotrophic larvae, and in the swimming lecithotrophic larvae; endodermal cells forming circular spots at the mid and lower trunk and in the swimming chaetal sacs in the 3-eye stage of the planktotrophic larvae and in the early-planktotrophic larvae; endodermal cells forming circular spots unevenly distributed in segments 8-13 in the late-planktotrophic larvae, in segments 9-12 in the planktotrophic juvenile and lecithotrophic juvenile. *Post1* is expressed earlier in the planktotrophic larvae than in the lecithotrophic larvae (2-eye stage for planktotrophic larvae and 3-eye stage for lecithotrophic larvae).

**Fig. 4A.**
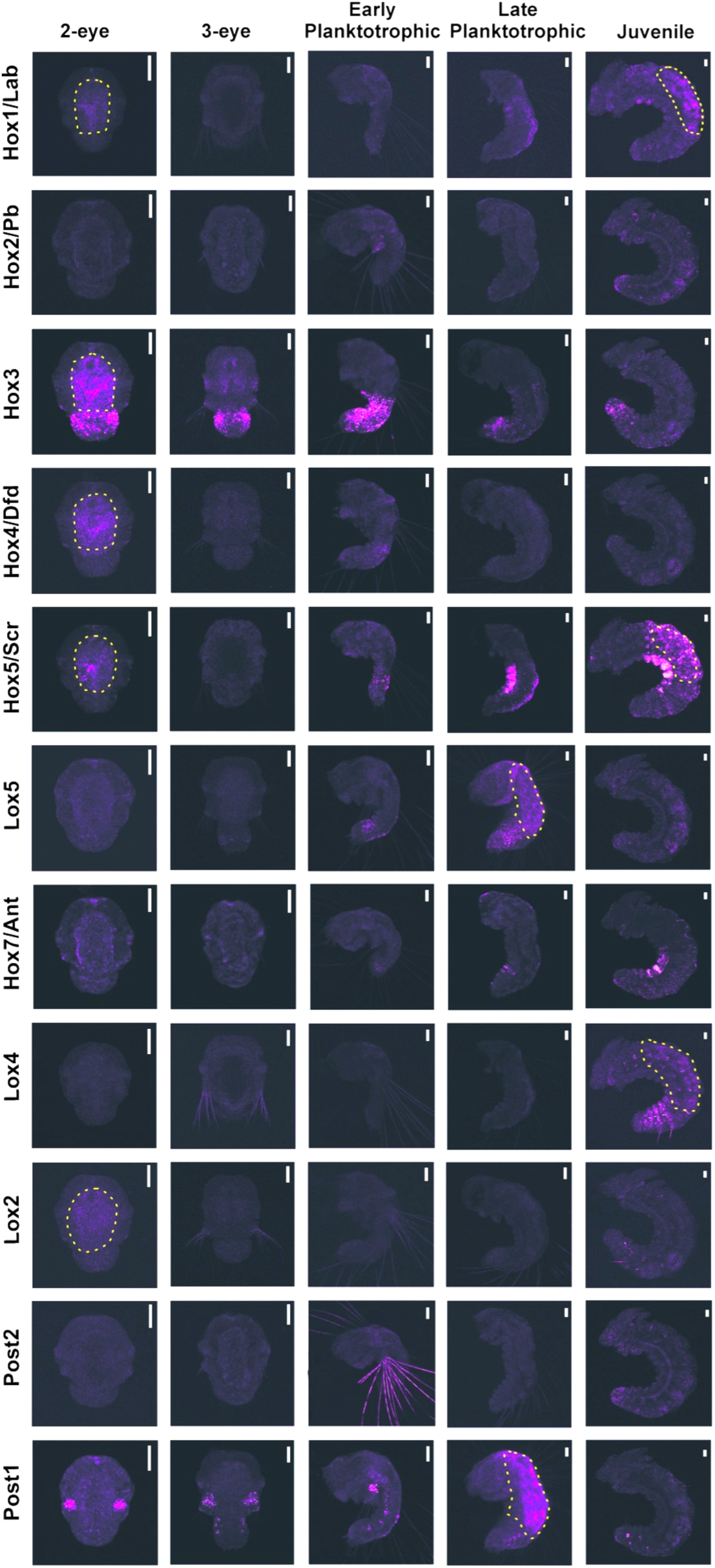
HCR *in situ* hybridization of the *S. benedicti* Hox genes at distinct developmental stages of the planktotrophic developmental mode. Yellow dashed lines encircle confirmed autofluorescent regions. Scale bars= 50 µm.

**Fig. 4B.**
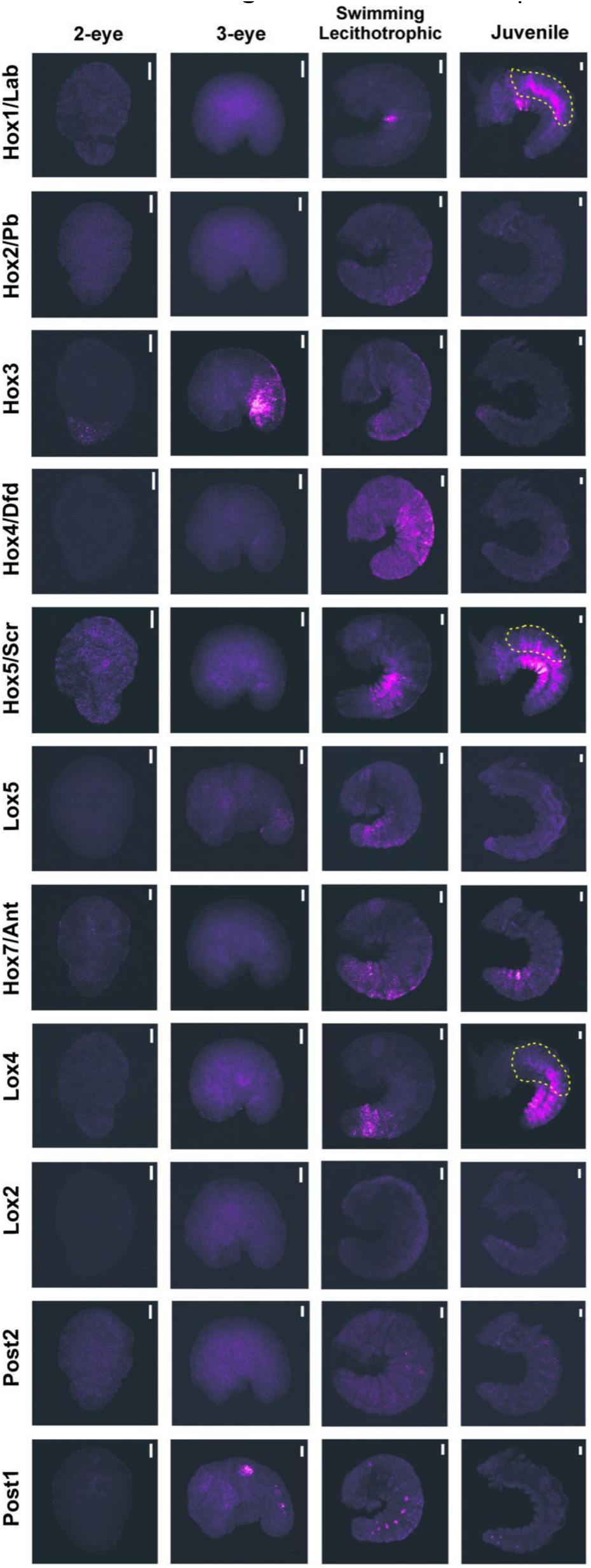
HCR *in situ* hybridization of the *S. benedicti* Hox genes at distinct early developmental stages of the lecithotrophic developmental mode. Yellow dashed lines encircle confirmed autofluorescent regions. Scale bars= 50 µm.

### Hox expression between the developmental morphs

*Heterochronies (change in timing):* Some Hox genes have a heterochronic shift in the timing of their expression. For *S. benedicti*, *Lox4* is expressed earlier in the swimming lecithotrophic larval stage than in the planktotrophic juvenile stage. But it has the same relative location of expression. *Post1* is expressed earlier in the 2-eye planktotrophic larval stage than in the 3-eye lecithotrophic larval stage (Figs. 4A,4B,6). *Dfd* and *Pb* stop being expressed one stage earlier in planktotrophic larvae (they disappear by the late-planktotrophic stage despite being present in the swimming lecithotrophic larvae).

*Heterotopies (change in location):* In the first chaetal sacs, which only form swimming chaetae in the planktotrophic larvae (and is a key morphological difference between the two developmental types), *Pb* and *Post1* are present in only the early-planktotrophic larvae. *Pb* has a different location in the swimming lecithotrophic larvae: cells in the latero/ventral part of the upper trunk (Fig. S2). *Post1* is also spatially different between the developmental types, appearing in the swimming chaetal sacs of the early-planktotrophic larvae and not the lecithotrophic larvae. In other stages it is expressed similarly across the two developmental modes. (Figs. 4A,4B,5,6).

**Fig. 5.**
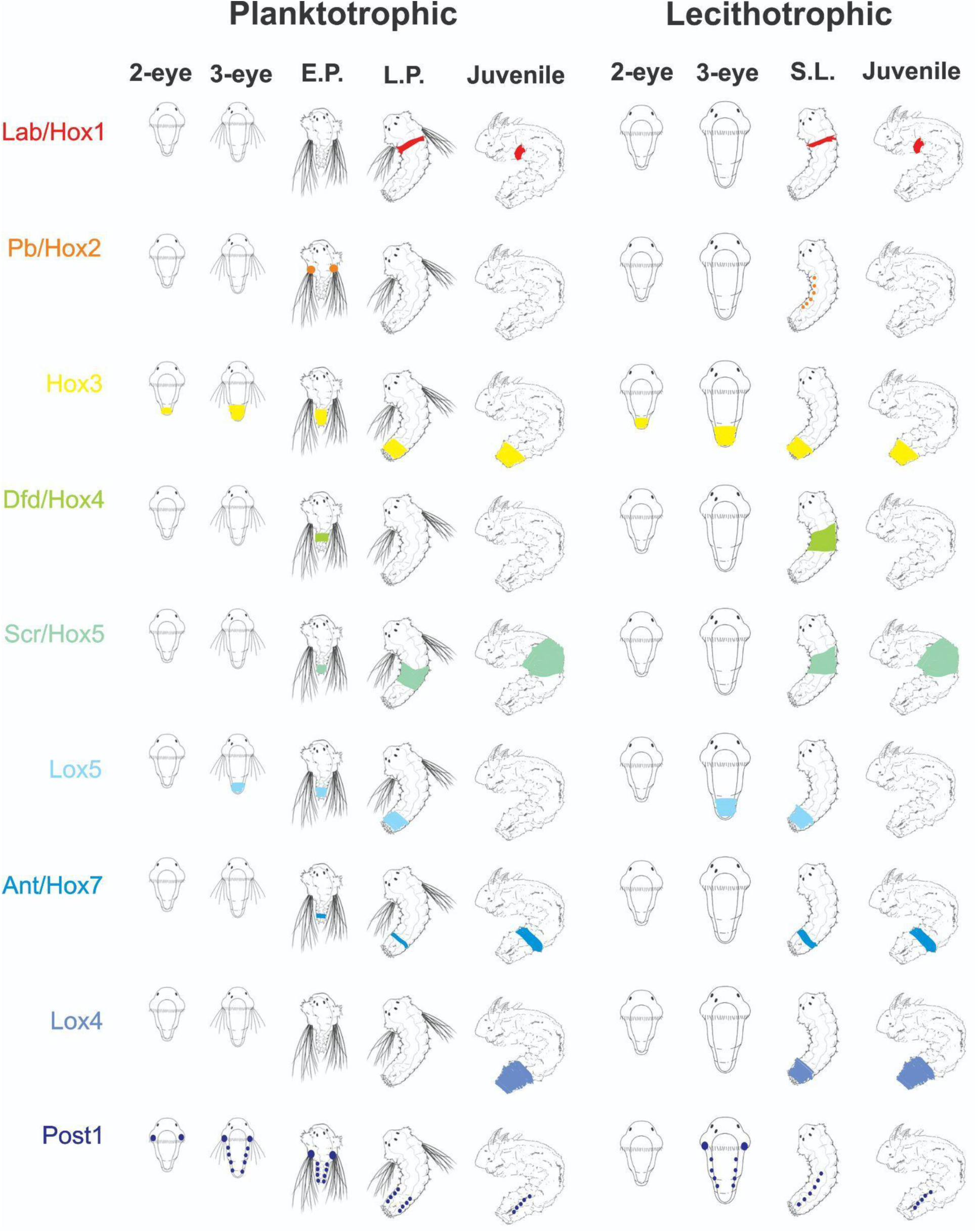
Schematic representation of the Hox gene expression patterns at distinct early life stages in the two developmental modes of *S. benedicti*. E.P.=early-planktotrophic, L.P.= late-planktotrophic, S.L.= Swimming Lecithotrophic.

**Fig. 6.**
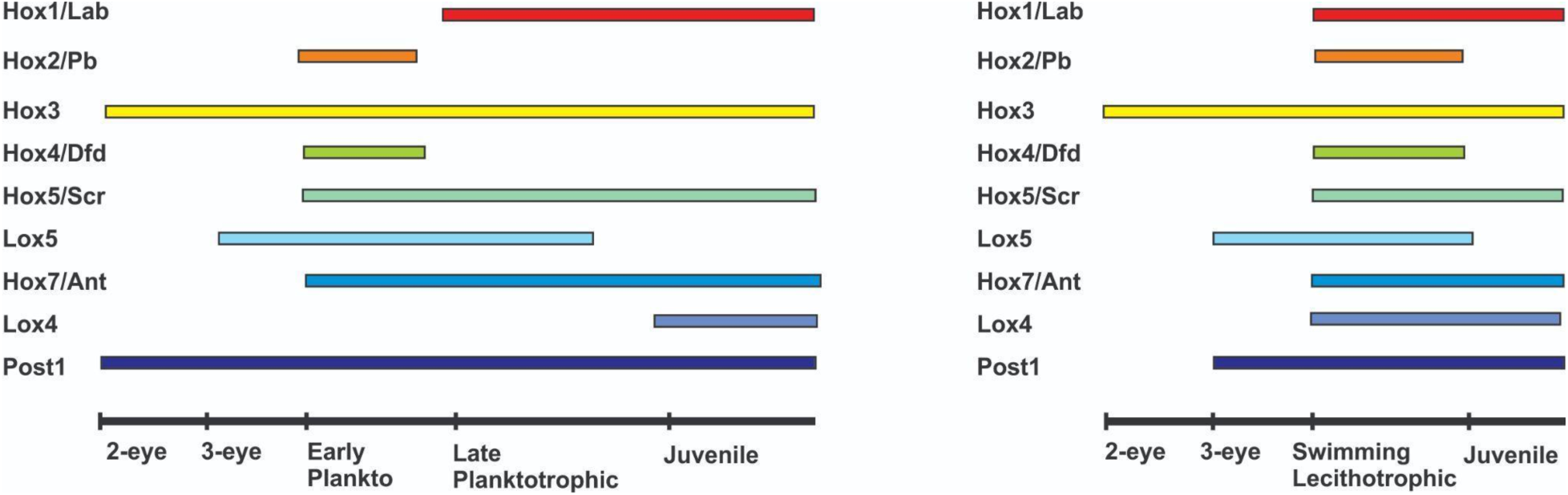
Timing of Hox gene expression in the two developmental modes of *S. benedicti*. Planktotrophic developmental mode is left and lecithotrophic developmental mode is right.

### RNA expression levels of the Hox genes

While we can visualize the onset and expression levels of the Hox genes using HCR *in situ* hybridization, we also wanted to compare Hox gene expression using a RNAseq approach. We sequenced RNA from three developmental stages across both types: 16-cell, blastula, gastrula, 2-eye larvae, swimming larvae which are ∼5-7 dpf (early-planktotrophic larvae and early swimming lecithotrophic larvae that we manually removed from the mother’s brood pouch), and 1-week old larvae which are seven dpf (‘late-planktotrophic’ larvae and ‘swimming lecithotrophic’ larvae; Harry and Zakas, 2023; Figs. 1, 7). We used RNAseq data both to identify Hox mRNAs for the HCR probe design and to quantify differential gene expression. We cannot detect any gene expression in these first three stages when using HCR, but we can compare the transcriptomic reads to the HCR patterns for the last three stages (Fig. 7). Although it is important to note that RNAseq and HCR are fundamentally different methodologies, and while patterns may be similar, we do not expect the results to directly translate across datasets.

**Fig. 7.**
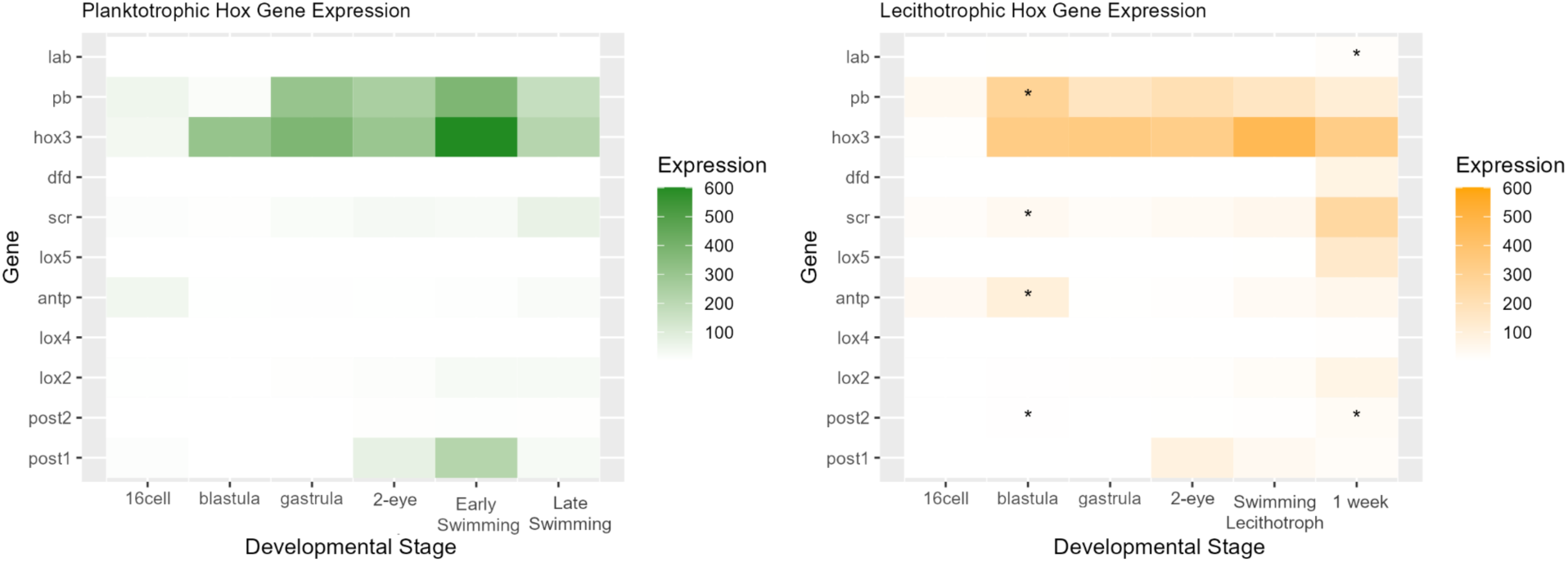
Heatmap of RNAseq Hox gene expression. Number of reads mapped to each gene over development shows similarity in Hox gene expression patterns across the types. Asterix indicates the gene is significantly differentially expressed at that time point compared to the other larval type as determined using DESeq2 and the full transcriptome dataset to assign significance. The six timepoints are equivalent stages across the two larval types. “1 week” larvae have not metamorphosed into juveniles and are late-stage swimming larvae.

Overall Hox gene expression is quite low in the RNAseq dataset, but very similar between morphs. Some statistically significant differences in expression are indicated (based on the criteria for a minimum fold-change difference in expression greater than 2, and a *p <0.05* using the entire transcriptome with DEseq). Most significant differences in expression are at the blastula stage, with later stages having largely conserved Hox gene expression. This is consistent with other spiralian patterns of gene expression, where typically there is more divergence (across species) or variation (within species) in gene expression in early embryogenesis that converge around gastrulation (Martín-Zamora et al. 2023; Harry and Zakas, 2023).

## DISCUSSION

This is the first comparison of developmental divergence in Hox genes expression within a species, which demonstrates the extent to which Hox genes underlie the developmental differences in larvae that make up alternate life-history modes. However, we see little evidence that the Hox genes drive developmental differences or are responsible for the divergence we see in overall larval morphology. Likely, the life-history differences and larval morphology arise from genes downstream of Hox expression in the developmental program.

### *S. benedicti* Hox homolog conservation with other annelids

The eleven Hox genes we identified, and their order in the genome, are homologous to other spiralians, particularly other annelids (Kourakis and Martindale, 2001; Peterson et al., 2000; Endo et al., 2016; Fröbius et al., 2008; Bakalenko et al., 2013; Li et al., 2020; Martin-Zamora et al., 2023). The split anterior-posterior clustering occurs in other annelids, brachiopods and mollusks (Fröbius et al. 2008; Schiemann et al., 2017; Martin-Zamora et al. 2023). The most closely related species where the Hox genes have been studied, the lecithotrophic annelid *Capitella teleta,* has a similar genomic organization and expression patterns in the swimming lecithotrophic larval type (Fröbius et al., 2008, ref for relatedness). In both *C. teleta* and *S. benedicti,* genes *Dfd* and *Lox5* are expressed in the mid and lower trunk of the segmented body, while *Scr* in the mid trunk and *Lox4* in the lower trunk (Fröbius et al., 2008).

Most bilaterians including the annelids *C. teleta*, *O. fusiformis*, *A. virens*, *N. virens* and *P. dumerilii,* generally follows the expected Hox pattern of spatial and temporal collinearity in expression over developmental time and segment addition (Kulakova et al., 2007; Bakalenko et al., 2013; Martin-Zamora et al., 2023). In *S. benedicti* we generally see the same pattern of spatial collinearity, with some expectations noted below, but we do not see the same pattern of temporal collinearity.

Temporal disruption could be consistent with the need to maintain variable developmental programs in a single species. One explanation for the early expression of more ‘anterior’ Hox genes is secondary co-option of gene function. Canonical Hox gene function is the specification of anterior-posterior body segments, but non-canonical functions can arise leading to secondary expression patterns in different locations or times (Hiebert and Maslakova, 2015; Schiemann et al., 2017; Fröbius and Funch, 2017). For example, temporal changes in both *S. benedicti* larvae compared to other bilaterians include early expression of *Hox3* and *Post1* (in both HCR and RNAseq data). *Post1* and *Pb* are expressed in the chaetal sacs leading to the possibility of earlier expression in these locations as a secondary co-option into a role in chaetogenesis. Similar atypical temporal patterns are also observed in some other annelids: *Hox3* is also expressed early in *C. teleta* (in RNAseq data; Martin-Zamora et al., 2023); In *P. duremilli, Pb* is expressed early at the internal zone of the swimming chaetal sacs in the late-trochophore stage (Kulakova et al., 2007), supporting that it may have a role in chaetal specification or initiation in annelids. *Post1* is expressed very early in *Nereis virens* based on *in situ* hybridization, and has a role in chaetogenesis (Kulakova et al., 2002; 2007).

The spatial colinearity is generally what we expect for both larval types (Fig 7). However, some genes are expressed in places that are not consistent with segment identity. For example, *Post1* and *Pb* are expressed in chaetal sacs, which occurs in other annelids and brachiopods and indicates it could be a spiralian-specific Hox role for the formation of body chaetae (Kulakova et al., 2002; 2007; Schiemann et al., 2017). The disruption of spatial collinearity in the expression patterns of some Hox genes are observed in other spiralian species of annelids, mollusks, brachiopods, rotifers and nemertean (Fröbius and Funch, 2017; Fröbius et al., 2008; Hiebert and Maslakova, 2015; Schiemann et al., 2017; Salamanca-Diaz et al., 2021; Martin-Zamora et al. 2023).

No expression was detected for *Lox2* and *Post2* at any of the developmental stages of *S. benedicti*. *Post2* and *Lox2* have been reported as the most highly expressed Hox genes at the posterior end of the growing zone in the juveniles of other species of annelids such as: *C. teleta, P. dumerilli* and *O. fusiformis* (Kulakova et al., 2007; Fröbius et al., 2008; Martin-Zamora et al. 2023). Further HCRs in the late juvenile stage from the two developmental modes should be undertaken to corroborate whether *Lox2* and *Post2* are expressed later at the posterior end of the growth zone.

### Hox expression between the developmental morphs within *S. benedicti*

Despite the developmental and life-history differences between the larval modes, we see remarkably little differentiation between Hox gene expression in either the HCR or RNAseq datasets. For both developmental modes, the earliest Hox genes expression is at the 2-eye stage when the body segments just begin to arise. Most Hox genes turn on at stages when the segments (trunk) are beginning to form (3-eye stage and swimming larval stage). The similarity in expression pattern between the larval types highlights that Hox gene expression, and more broadly its relationship to body plan specification and initiation, are not different between animals based on their trophic life-history mode. Rather, the differences in Hox timing we see across other spiralian species with alternate life-histories are likely due to larger phylogenetic and evolutionary differences.

While the majority of Hox gene expression is similar between the two developmental modes, there are some key differences. Notably, there are heterotopic and heteromorphic changes in some Hox gene expression. Heterotopic differences in expression occur in *Pb* and *Post1,* which may have been secondarily co-opted in planktotrophic larvae to regulate swimming chaetae formation between the two developmental modes. Heterochronies across species are difficult to detect due to the methodological barriers of assigning equivalent stages across divergent species. Changes in the relative timing of expression could lead to segment specification differences and drive morphological change across lineages (Onimaru et al., 2021; Drobreva et al., 2022). In *S. benedicti*, we see clear patterns of heterochrony in Hox gene expression across the types at relative stages (*Lox4* and *Post1*), indicating that changes in the regulatory timing may be easily evolved. However, It is unlikely that the heterochronic shifts we see in two Hox genes are a major driver of body plan and life-history diversification within *S. benedicti*, although they could contribute to specific morphological differences between the two types like the formation of swimming chaetae.

Despite being a single species, assigning equivalent stages within *S. benedicti* is also difficult at some stages. Early embryological timepoints are clearly morphologically equivalent, but the larval stages are more difficult to compare. For example, the stage of larval release (‘early-planktotrophic’ larvae and ‘swimming lecithotrophic’ larvae) are equivalent stages in terms of many life-history traits. However, morphologically the ‘swimming lecithotrophic’ larvae are more similar to the ‘late-planktotrophic’ stage. Our Hox gene analysis clearly demonstrates that in terms of body plan formation, the former stages (‘early-planktotrophic’ larvae and ‘swimming lecithotrophic’ larvae) are the most equivalent despite the size and morphological differences. We have also recently completed an extensive gene expression analysis over larval development of the two morphs (Harry and Zakas, 2023), which confirms that these are equivalent stages. Because the planktotrophic larvae have a longer and more differentiated larval phase covering two time points, where the lecithotrophic larvae have one, it leads to destinations of minor heterochronies. Namely, it is this staging difference that highlights the slightly different timing of *Lab*, *Pb* and *Dfd* between the morphs.

When addressing heterochronies, it is critical to note that we are comparing the timing of gene expression at equivalent stages that differ in absolute time. In this way, the small number of heterochronic shifts we see are more surprising: consider that to have gene activation at the same relative time in development, the lecithotrophic offspring must initiate expression later in real time with a larger embryo (and cell) size than the planktotrophic larvae. The mechanisms that keep the same relative timing of expression in embryos of different sizes and ages are unknown. One theory of gene expression regulation in early embryos involves the nucleocytoplasmic (N:C) ratio (the proportion of cytoplasm to nuclear volume mediates transcriptional timing; Wolf, 2009). But this same N:C ratio would occur at different times in the two developmental types. Absolute time, or maternal timing, is also a key regulator in terms of transcript degradation or molecular half-lives (Buccitelli and Selbach, 2020; Lee et al., 2020); maternal (or zygotic) transcript degradation may occur at a different stage in the planktotrophic larvae compared to the lecithotrophic larvae. Again, a simple increase in egg size in the lecithotrophic type is not necessarily sufficient to explain the differences and similarities we see between the two developmental types.

Hox genes in *S. benedicti* are not a major contributor to morphological and life-history differences at the level of body plan. While overall patterns of Hox gene timing and location are quite similar in both types in the HCR and RNAseq data, there are multiple instances where Hox genes can be contributing to key life-history differences: some Hox genes (*Pb* and *Post1*) are clearly expressed differently in chaetal sacs and could have undergone a secondary co-option to regulate the formation of swimming chaetae in planktotrophic larvae. There are few heterochronic shifts in expression between the equivalent stages of the two developmental morphs, but whether these changes initiate any significant timing of downstream developmental differences remains to be determined.

## MATERIALS AND METHODS

### Identification of the Hox Genes in the genome and transcriptome

RNAseq reads from six developmental stages (Harry and Zakas, 2023) of both planktotrophic and lecithotrophic offspring were assembled with long-read (iso-seq CCS reads) guidance using the Trinity assembler (Grabherr et al., 2011). Hox gene transcripts were identified from the resulting contigs using tblastn (e-value cutoff 1*10^-30) from the BLAST command line tools (Altschul et al., 1990) and a set of Hox gene queries taken from other related taxa/species (Supplementary DataSheet). Eleven different contigs corresponding to eleven paralog genes were located in the genome of *S. benedicti* (Zakas et al., 2022) by using JBrowse (Skinner et al., 2009). The genomic location and gene length for each Hox gene is characterized in Table S3.

### Molecular Phylogenetic Analyses

We used Hox genes of other spiralian species from Genbank: *Platynereis dumerilii* (Kukalova et al., 2007; Maslakov et al., 2021), *Capitella teleta* (Frobius et al., 2008), *Owenia fusiformis* (Martin-Zamora et al., 2023), *Crassostrea virginica* (Modak et al., 2021) and *Lottia goshimai* (Huan et al., 2020) to construct a gene phylogeny (Table S2, Supplementary Data1). Amino acid sequences were aligned using MUSCLE which is included in the SEAVIEW software (Edgar, 2004; Gouy et al., 2010). A conservative alignment strategy was employed where all the positions that were spuriously aligned were excluded. The final alignment contains 64 sequences with 264 amino acid sites (Supplementary Data 2). Maximum-likelihood phylogenetic trees were constructed using the IQ-Tree software (Nguyen et al., 2015). VT+I+G4+F was the best fit evolutionary model and - 28796.8292 was the optimal log likelihood. Support values of the ML tree were calculated by three different methods: 1000 ultrafast bootstrap replicates (Minh et al., 2013), 1000 replicates of the Shimodaira–Hasegawa approximate likelihood ratio test (SH-aLRT), and an approximate eBayes test (Anisimova et al., 2011) (Figure 3, S1).

### Animal Culturing

We collected embryos and larvae at each stage from lab reared animals of each developmental type as in Zakas (2022). Lecithotrophic worms were collected from Long Beach (California) and planktotrophic worms from Newark Bay (New Jersey; Zakas et al., 2018).

### Hybridization Chain Reaction (HCR)

Multiplex probes of the eleven Hox gene transcripts of *S. benedicti* were designed using the HCR3.0 Probe Maker (Kuehn et al., 2022) (Table S1). Exact oligo sequences and the associated hairpins are listed in the Supplementary DataSheet. The sequences generated by the software were used to order different batched DNA oligo pools (50 pmol DNA Pools Oligo Pool) from Integrated DNA Technologies, resuspended to 1 pmol/μl in Nuclease Free Water. Different developmental stages from the two modes of *S. benedicti* were fixed in 4% paraformaldehyde at 4°C overnight, transferred stepwise into 100% methanol, and kept at −20°C. Samples were rehydrated through a methanol/DEPC-treated PBSt series. Probe hybridization buffer, probe wash buffer, amplification buffers, and a DNA HCR amplifier hairpin set were purchased from Molecular Instruments. HCR was performed as previously described (Choi et al., 2018; Kuehn et al., 2022).

### Imaging

HCR samples were mounted in Slowfade Glass with DAPI and kept at 4°C until imaging and imaged using Zeiss Laser Scanning Confocal Microscope LSM 710. Z-stack images (32 layers) were processed in ImageJ (Abràmoff et al., 2004). At least two replicates for each sample were imaged to corroborate that the gene expression patterns detected were the same in different samples. HCR probes were multiplexed so that imaging a single individual would capture expression for 2-3 hox genes (Table S1). Gene expression was considered real (and not background fluorescence) when a region only showed expression under one of the three possible color channels with distinct fluorescent spectra. (Table S1). We considered regions with fluorescence under multiple channels to be autofluorescence in the sample. In addition, samples from different Hox genes that have the same channel (same hairpins) were compared to identify any autofluorescence (Figs. S3 and S4). Control animals with only hairpins and no probes were also used to determine regions with background fluorescence. Chitinous regions of the larvae autofluoresce in some channels (mainly the green wavelength 488; as in the swimming chaeate of Fig. 4A early-planktotrophic larvae *Post2).* Endodermal expression was considered as an intense fluorescence of the first plane from the Z-stack images opposite to the last plane that it could be considered ectodermal.

### RNA seq data

Hox gene expression data was generated using RNAseq (Harry and Zakas, 2023). Each developmental stage consisted of three to five biological replicates. Raw RNAseq reads were quality trimmed using FastP (Chen et al., 2018) and TrimGalore (cutadapt) (Martin, 2011) and then mapped to a reference transcriptome using Salmon (Patro et al., 2017). The reference transcriptome was assembled using RNAseq reads and previously published IsoSeq data (Zakas et al 2022) in conjunction with the Trinity assembler (Grabherr et al., 2011) and assembled transcripts derived from Hox genes were identified using BLAST (Altschul et al., 1990). Sample expression quantification estimates were then normalized and statistical tests for differential expression between morphs at each stage were performed using the standard DESeq (Love et al., 2014) workflow for time-series experiments in R. Heatmaps were made in R using the ggplot2 package (Wickham, 2016).

## Acknowledgements

Kayleigh McHugh helped with animal collection, maintenance and fixation of samples. Mariusz Zareba assisted with the confocal analysis. Duygu Özpolat, Bria Metzger, Emma Kelley, and Veronica Acosta assisted with troubleshooting HCR *in situs*, and Ryan Null developed the probe finder for constructing HCR probes. Bruno Pernet, Carrie Albertin and Matthew Rockman provided comments on the manuscript.

## Funding Information

The initial work was completed at the Marine Biological Labs, Woods Hole MA with a Whitman Fellowship granted to. Zakas. This work is supported by the NIH MIGMS grant 5R35GM142853 to C. Zakas.

## Data Availability

RNA reads are available at NCBI under BioProject PRJNA1008044.

## Competing interests

The authors declare no competing or financial interests.

## SUPPLEMENT

**Fig. S1.**
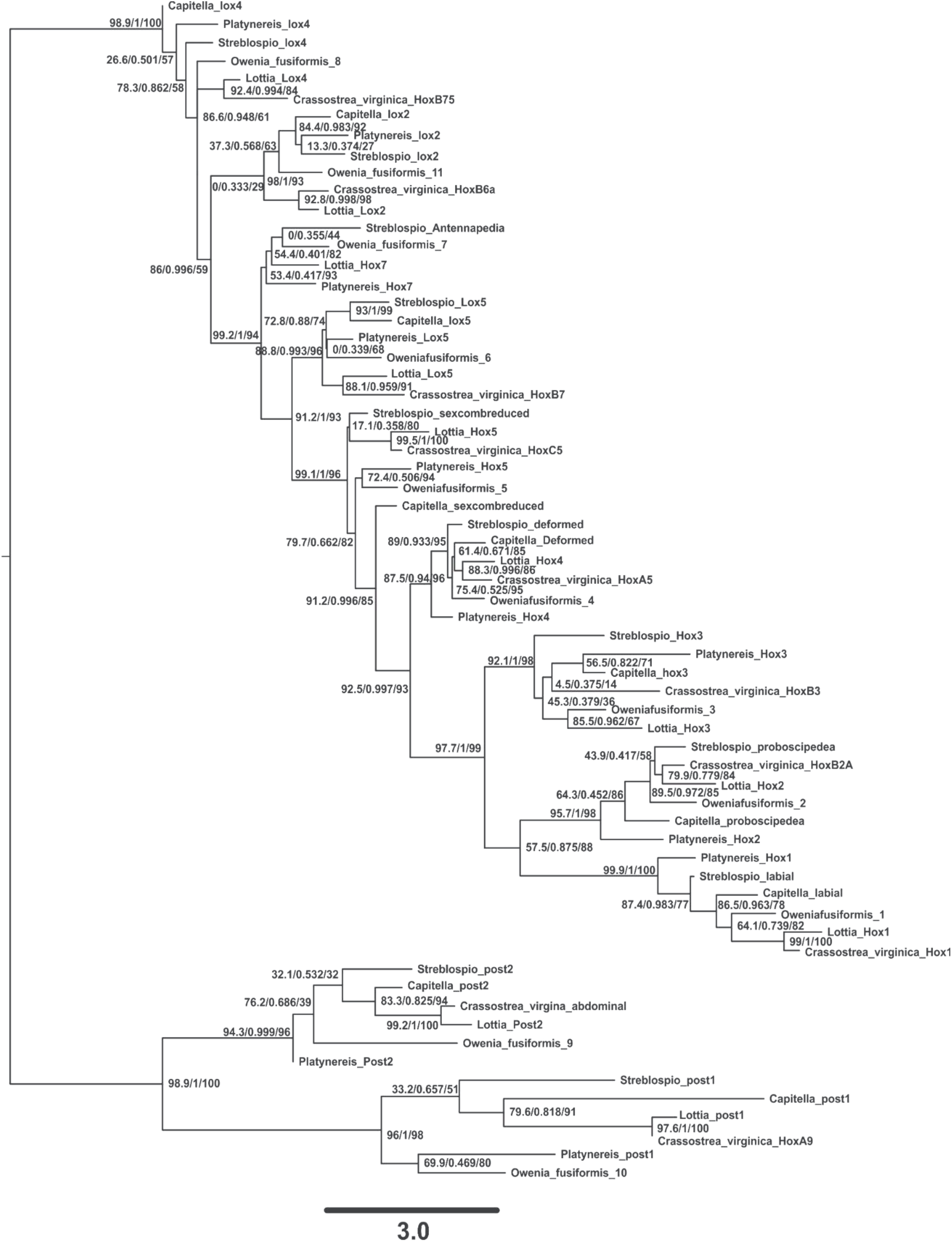
Molecular phylogenetic tree of Spiralian Hox genes. The tree was constructed with the IQ-Tree software. Bootstrap values for the three different methods (see Material and Methods) calculated in IQTREE software are shown in the branches.

**Fig S2.**
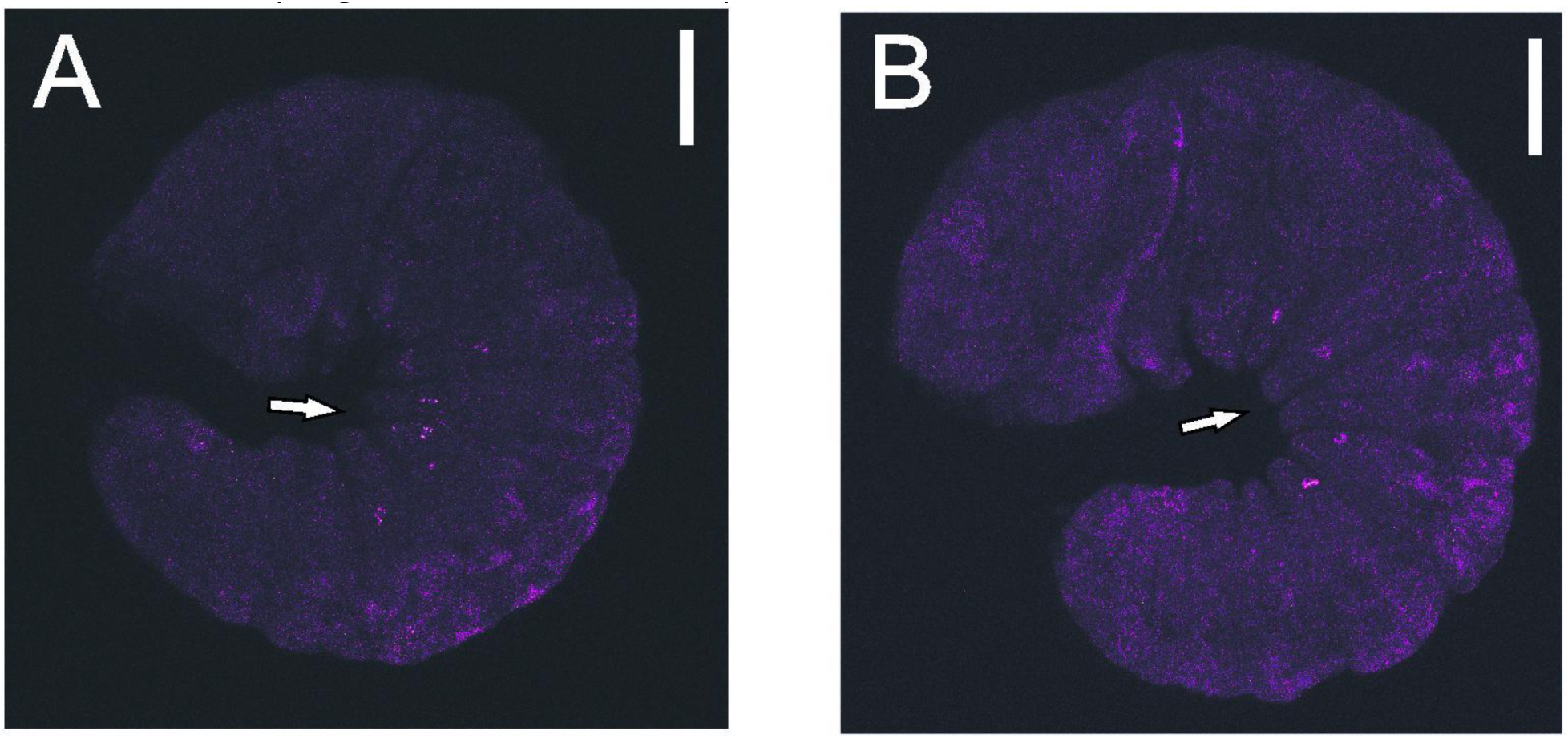
HCR *in situ* hybridization of *Pb* at the swimming larval stage in the lecithotrophic larvae. The arrows indicate the cells expressing the gene that form circular spots at the latero-ventral area in each body segment. Scale bars= 120 µm.

**Fig S3.**
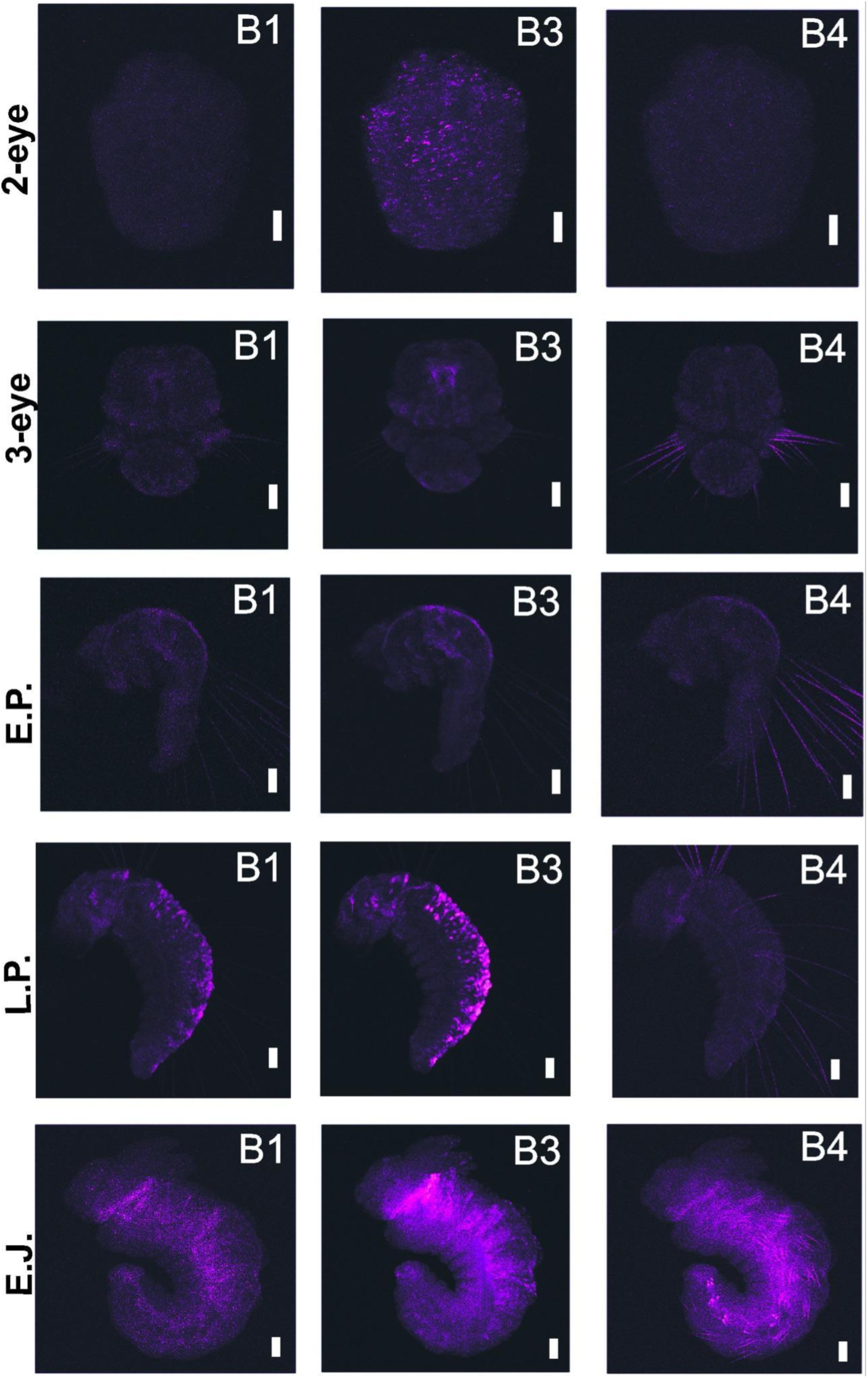
HCR *in situ* hybridization of the control samples of the planktotrophic larvae at distinct developmental stages. We consider all fluorescence captured without probes as background. Fluorescent spectrum of each channel is: B1-594, B3-647 and B4-488. E.P.=early-planktotrophic, L.P.= late-planktotrophic and E.J.= Juvenile. Scale bars= 50 µm.

**Fig S4.**
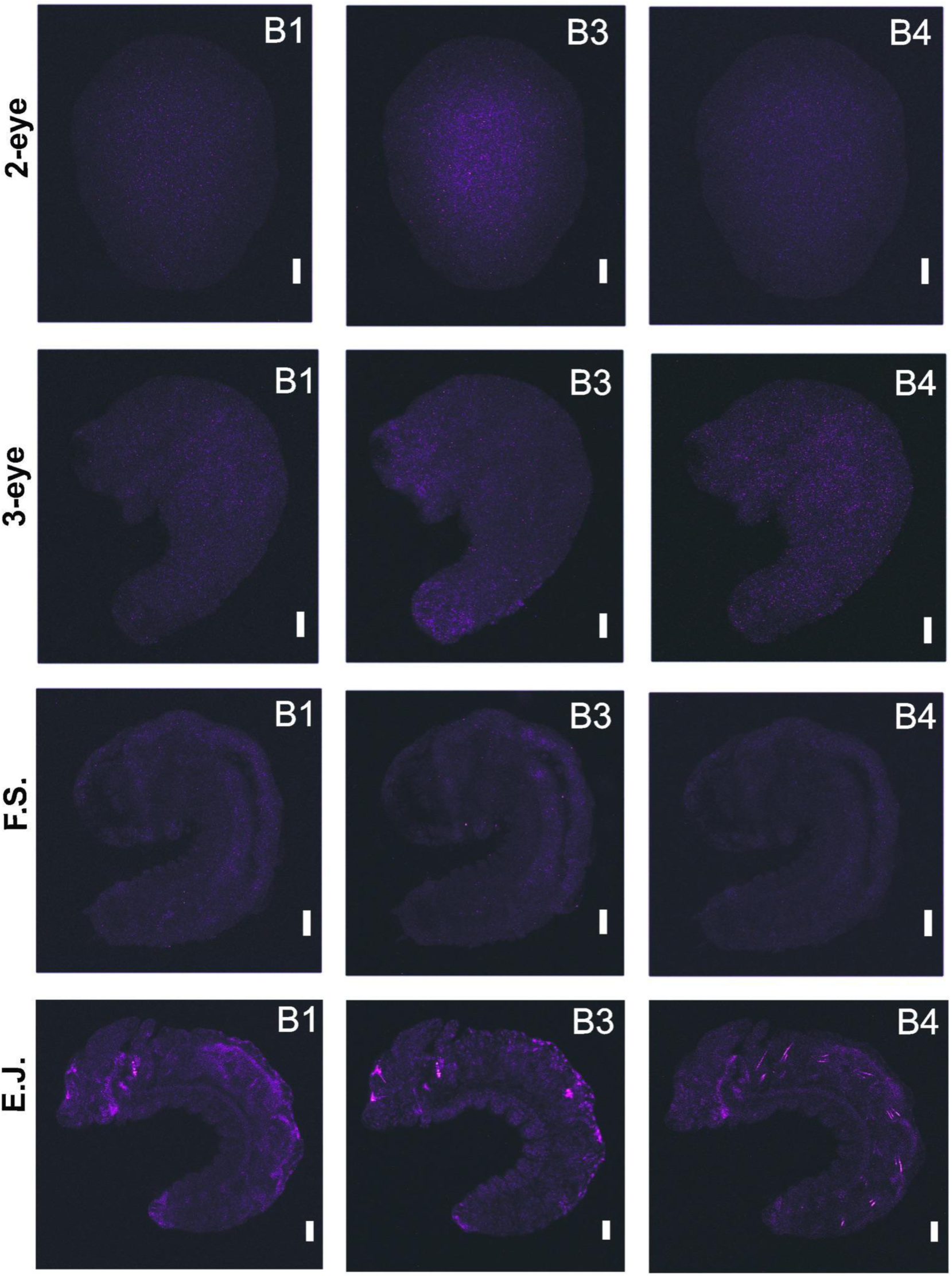
HCR *in situ* hybridization of the control samples of the lecithotrophic larvae at distinct developmental stages. We consider all fluorescence captured without probes as background. Fluorescent spectrum of each channel is: B1-594, B3-647 and B4-488. F.S.= Swimming Stage and E.J.= Juvenile. Scale bars= 50 µm.

**Table S1.** Set of groups of the eleven Hox genes that were used to carry out the multiplexed HCR *in situ* hybridization experiment to identify their expression patterns. B1, B3 and B4 are the HCR hairpins. Fluorescent spectrum of each channel is: B1-594, B3-647 and B4-488.

| Group1 | Group2 | Group3 | Group4 |
| --- | --- | --- | --- |
| Lab-B1 | Pb-B1 | Lox5-B1 | Dfd-B1 |
| Scr-B3 | Ant-B3 | Post1-B3 | Hox3-B3 |
| Lox4-B4 | Post2-B4 |  | Lox2-B4 |

**Table S2.** Genbank accession numbers of the Hox gene sequences from spiralian species that were used to construct the molecular phylogenetic tree. Pd= *Platynereis dumerilii,* Ct= *Capitela teleta*, Of= *Owenia fusiformis*, Cv= *Crassostrea virginica* and Lg=*Lottia goshimai.* * The sequences of Ant and Lox4 from *Crassostrea virginica* are the same. The sequences of *Lox4* and *Lox2* from *P. dumerilii* were obtained from an assembled transcriptome (Maslakov et al., 2021). The sequence of Ant from *Capitela teleta* was too short and not included in the molecular phylogenetic tree.

|  | Pd | Ct | Of | Cv | Lg |
| --- | --- | --- | --- | --- | --- |
| Hox1/Lab | AFJ91921.1 | ABY67952.1 | CAH1773442.1 | XP_022291943.1 | QCF47216.1 |
| Hox2/Pb | AFJ91922.1 | ABY67953.1 | CAH1773441.1 | XP_022289925.1 | QCF47217.1 |
| Hox3 | AFJ91923.1 | ABY67954.1 | CAH1773440.1 | XP_022292027.1 | QCF47218.1 |
| Hox4/Dfd | AFJ91924.1 | ABY67955.1 | CAH1773439.1 | XP_022307961.1 | QCF47219.1 |
| Hox5/Scr | ATG29892.1 | ABY67956.1 | CAH1773438.1 | XP_022340779.1 | QCF47220.1 |
| Lox5 | AFJ91925.1 | ABY67957.1 | CAH1773437.1 | XP_022308431.1 | QCF47221.1 |
| Hox7/Ant | ATG29893.1 | ABY67962.1 | CAH1773436.1 | * | QCF47222.1 |
| Lox4 |  | ABY67958.1 | CAH1773435.1 | XP_022342129.1 | QCF47223.1 |
| Lox2 |  | ABY67959.1 | CAH1773434.1 | XP_011455564.1 | QCF47224.1 |
| Post1 | AFJ91928.1 | ABY67961.1 | CAH1775937.1 | XP_022291104.1 | QCF47226.1 |
| Post2 | AFJ91927.1 | ABY67960.1 | CAH1773433.1 | XP_022286889.1 | QCF47225.1 |

**Table S3.** Table showing the eleven hox genes with their genomic location, transcript length, gene length and the number of probes that were designed for the HCR experiment.

| Gene | Genomic Location | Transcript Length (bp) | Transcript bases used in HCR probes (bp) | Number of probes | Gene Length (bp) |
| --- | --- | --- | --- | --- | --- |
| lab | 6119808-6120980 (CH 7) | 2228 | 1,305 | 29 | 1172 |
| pb | 6093553-6096033 (CH 7) | 3618 | 1,800 | 40 | 2480 |
| hox3 | 6060233-6063423 (CH 7) | 4778 | 2,115 | 47 | 3190 |
| dfd | 5955740-5956747 (CH 7) | 784 | 585 | 13 | 1007 |
| scr | 5890873-5894127 (CH 7) | 4234 | 1,800 | 40 | 3254 |
| lox5 | 5845156-5846927 (CH 7) | 3462 | 1,800 | 40 | 1771 |
| antp | 5799538-5801770 (CH 7) | 2477 | 1,350 | 30 | 2232 |
| lox4 | 5658073-5658834 (CH 7) | 3639 | 1,800 | 40 | 761 |
| lox2 | 45895179-45898552 (CH 7) | 3382 | 1,665 | 37 | 3373 |
| post2 | 45865494-45866205 (CH 7) | 1392 | 810 | 18 | 711 |
| post1 | 12597997-12599246 (CH 7) | 2338 | 1,350 | 30 | 1249 |

## LEGENDS OF THE SUPPLEMENTARY DATASETS

**Supplementary Data1** – Amino acid sequences of the Hox genes of Spiralians that were used to build the molecular phylogenetic tree (no alignment).

**Supplementary Data2**-Amino acid sequence regions of the Hox genes that were spuriously aligned to build the molecular phylogenetic tree.

**Supplementary Data3**

**Sheet1 -** Table containing sequences of the hairpins designed and synthesized of the eleven Hox genes of *S. benedicti*.

**Sheet2 -** Table containing the characteristics of the BLAST searches for the identification of the eleven Hox gene sequences from the transcriptome of *S. benedicti*.

**Sheet 3 -** Tables containing the number of read counts for selected developmental stages in the two modes of *S. benedicti* from the transcriptomic datasets analyses.

## Notes

### Competing Interest Statement

The authors have declared no competing interest.

